# Nanoparticle Tracking Analysis: A powerful tool for characterizing magnetosome preparations

**DOI:** 10.1101/2020.06.23.166587

**Authors:** Alfred Fernández-Castané, Hong Li, Stephan Joseph, Moritz Ebeler, Matthias Franzreb, Daniel G. Bracewell, Tim W. Overton, Owen R.T. Thomas

## Abstract

Nanoparticle Tracking Analysis (NTA) has been employed to measure the particle concentration and size distribution of magnetosomes extracted and purified from *Magnetospirillum gryphiswaldense* MSR-1, and then exposed to probe ultrasonication for various times, or 1% (w/v) sodium dodecyl sulphate (SDS) for 1 h. Particle concentration increased 3.7-fold over the first 15 min of ultrasonication (from 2 × 10^8^ to >7.3 × 10^8^ particles mL^−1^), but fell steeply to ~3.6 × 10^8^ particles mL^−1^ after 20 min. NTA of untreated magnetosome preparation confirmed a wide particle distribution dominated by larger species (D[1,0] = 312 nm; D_n50_ = 261 nm; mode = 243 nm) with no particles in the size range of isolated single magnetosomes. After 5 min of ultrasonication the whole particle size distribution shifted to smaller size (D[1,0] = 133 nm; D_n50_ = 99 nm; mode = 36 nm, corresponding to individual magnetosomes), but longer treatment times (15 and 20 min) reversed the previous transition; all characteristic numbers of the particle size distributions increased and very few small particles were detected. Side-by-side comparison of NTA and TEM sizing data revealed remarkable similarity at low ultrasonication times, with both showing single magnetosomes accounted for ~30% population after 5 min. Exposure of magnetosomes to SDS resulted in a ~3-fold increase in particle concentration to 5.8 × 10^8^ particles mL^−1^, narrowing of the size distribution and gross elimination of particles below 60 nm. We conclude that NTA is a rapid cost-effective technique for measuring particle number, size distribution and aggregation state of magnetosomes in solution, but requires further work to improve its resolving power.

## 1. Introduction

Magnetosomes are functional magnetic nanoparticles that are naturally generated by magnetotactic bacteria (MTB) [1]. Magnetosomes are arranged as highly ordered chains of single-domain magnetite (Fe_3_O_4_) or greigite (Fe_3_S_4_) crystals individually coated in a phospholipid membrane and usually attached to a protein filament that aligns them with the axis of the bacterium [2, 3]. Composition not withstanding individual magnetosomes possess the key advantages of high ferrimagnetism, narrow size distribution (20–120 nm) and shape distribution [2–6] and biologically compatible surface chemistry [5, 7]. Furthermore, their membrane contains specific transmembrane proteins that can be exploited for biotechnological applications, e.g. as anchors for chemical and/or biochemical modification [8 – 11].

Despite numerous efforts to improve the yields of magnetosomes [4, 12, 13], their large-scale biomanufacture remains significant barrier to future widespread industrial application. Fundamental to this are: (i) appropriate means for analysing MTB growth and physiology [14, 15]; (ii) investigating the biomineralization of magnetic iron-containing minerals, which is still not fully understood [16 – 18]; (iii) the ability to both produce batches of nanoparticles that are consistent in structure, chain length and functionality – this is key to unlocking the potential of magnetosomes as therapeutic agents; and (iv) to be able to rapidly analyse these characteristics.

Transmission electron microscopy (TEM), the gold standard technique for characterization of MTB and magnetosomes since their discovery in the 1970s [1], is expensive, time consuming and importantly some of the information it conveys is not translatable to behaviour in solution. Most critically, the possible introduction of artefacts at many different points during sample preparation for TEM [19 – 21] can blur the state of the dispersion to the extent that number-weighted particle size distributions generated by automated image analysis cannot be trusted [21]. Other powerful techniques frequently employed in the chemical and magnetic characterization of magnetosomes include X-ray diffraction, Fourier-transform infrared spectroscopy, energy dispersive spectroscopy, Mössbauer spectroscopy, vibrating sample magnetometry, and alternating field gradient magnetometry [22 – 27].

With few exceptions the above methods, TEM included, do not satisfy the exacting criteria required for routine analytical tests in the development and quality control of magnetosomes as future nanomedicines, namely reproducibility, robustness, speed, and throughput. Crucial to magnetosome preparations are the number of magnetosome units per sample, given that this depends on chain length [15]. The relatively new technique of Nanoparticle Tracking Analysis (NTA) offers attractive prospects in this context, given the following:

i. its ability to directly visualize, size and count nanoparticles ‘particle-by-particle’ in liquid samples requiring no or limited pre-treatment;
ii. it provides accurate measures of hydrodynamic particle size in relatively rapid time and in cost effective manner;
iii. its suitability for real-time analysis of dilute polydisperse particle systems containing species ranging from a few tens on nanometers to 1 – 2 micron particles and reporting particle concentration in particles per millilitre [28, 29]; and
iv. impressive demonstrations of its utility with a diverse range of biological, synthetic and hybrid nanoparticles, including liposomes, exosomes, immunogenic complexes, drug delivery nanoparticles, biopharmaceutical protein aggregates, silica, metal and metal oxide nanoparticles, including magnetite [28 – 38].

Here we propose NTA as a future process analytical and quality control tool in magnetosome research and development. Specifically, we describe its use for characterizing changes in the particle size distribution of extracted magnetosome preparations following liquid shear and surfactant attack, processes known to degrade and destabilize chained magnetosome. We have correlated the observed changes in particle size distribution with TEM analysis, finding striking agreement between the two techniques at low ultrasonication times, and demonstrate that NTA’s use in detecting and studying aggregation of nanoparticles [32 – 36] is usefully extended to magnetosome preparations.

## 2. Materials and Methods

### 2.1. Strains, growth media and culture conditions

*Magnetospirillum gryphiswaldense* MSR-1 was obtained from Deutsche Sammlung von Mikroorganismen und Zellkulturen GmbH (DSMZ, Germany) and used for all experiments. MSR-1 cells were grown in bioreactors using pH-stat operation mode as described elsewhere [13]. Briefly, batch medium consisted of flask standard medium (FSM) [4] without iron citrate. The pH was controlled at a set-point of 7 with the automated addition of an acidic feeding solution, and cells were grown at 30 °C under microaerobic conditions, by maintaining pO_2_ <1% using a closed feedback loop regulating agitation between 100 – 500 rpm and manually supplying air between 0 – 100 mL min^−1^.

### 2.2. Cell harvesting and magnetosome purification

MSR-1 cells were harvested by centrifugation (7,500 g_av_, 20 min, 4°C) in a Beckman J2-21 centrifuge fitted with a fixed-angle JA-14 rotor (Beckman Instruments Inc., Palo Alto, CA, USA; Beckman Coulter Life Sciences, Indianapolis, IN, USA). The supernatant was removed and cells were stored at −18°C until further use. Subsequently, cells were thawed at 4°C and suspended in 50 mM HEPES buffer, pH 8.0, to a final wet cell concentration of 20% (w/v). Cells were disrupted in a single pass through a bench top high pressure homogenizer (Constant Systems Ltd., Daventry, Northants, UK) operated at 10 kpsi. The resulting homogenate was subjected to magnetic separation, and following resuspension and washing in phosphate buffered saline (PBS, pH 7.4), portions (1 mL) of the recovered magnetosomes were layered onto 4 mL 60% (w/v) sucrose cushions (prepared in 10 mM HEPES buffer, pH 8.0) contained in 10 mL Oak Ridge High-Speed PPCO screw-cap round-bottomed centrifuge tubes (Model 3119-0010, Thermo Fisher Scientific, Loughborough, Leics, UK). Samples were centrifuged at 50,000 g_av_ in the fixed angle rotor ‘model 12111’ (10 × 10 mL) of a Sigma 3K30 centrifuge (Sigma Laborzentrifugen GmbH, Osterode am Harz, Germany) for 2.5 h at 4°C. After centrifugation, the ‘light’ sucrose top phases were carefully removed using a Pasteur pipette, and the ‘heavy’ fraction at the bottom of each tube containing magnetosomes was collected and resuspended in PBS filtered through a 0.2 μm polyether sulfone syringe filter to a concentration of 35 or 70 mg iron L^−1^ (see section 2.4). The purified magnetosomes used in this work came from different fermentations, hereafter referred to as ‘batch 1’ (70 mg Fe iron L^−1^) and ‘batch 2’ (35 mg iron L^−1^).

### 2.3. Treatment of purified magnetosomes

Intentional alteration of the particle size distributions of purified magnetosomes was done in two different ways. In the first, 5 mL of a ‘batch 1’ magnetosomes diluted to 20 mL volume with PBS was subjected to ultrasonic disruption on ice using a Status US70 ultrasonic (20 kHz, 60 W) probe sonicator (Philip Harris Scientific, Lichfield, Staffs, UK) operated in 1 min bursts (50% duty cycle) at 70% amplitude (power) with 1 min of cooling of the probe in ice cold water between successive bursts. Five hundred microlitre samples were removed after various times (0, 1, 5, 15 and 20 min), brought to 22°C, passage through a 1 mL syringe fitted with a 29 gauge needle, and analysed in the LM10 (section 2.5) within 5 min of treatment. In the second, 5 mL portions of ‘batch 2’ magnetosomes were incubated in PBS supplemented with 1% (w/v) SDS or PBS alone. After 1 h at room temperature, 500 μL samples were withdrawn, passaged as indicated above and immediately analysed in the LM10 (section 2.5).

### 2.4. Determination of iron content

The iron concentration in samples was measured by atomic absorption spectroscopy as described previously [13] using a single element iron (248.3 nm) hollow cathode lamp (SMI-LabHut Ltd., Churcham, Glos, UK) operated at a current of 30 mA with an acetylene / air flame (0.7 L acetylene min^−1^ and 4.0 L air min^−1^) in a Perkin Elmer AAnalyst 300 Atomic Absorption Spectrometer (Waltham, MA, USA). Samples were prepared in triplicate. Briefly this involved solubilising the iron present with 500 μL of 70% (v/v) nitric acid and incubating at 98°C for 2 h with shaking at 300 rpm in an Eppendorf^®^ ThermoMixer^®^ Comfort 5355 dry block shaker (Eppendorf UK Ltd, Stevenage, Herts, UK).

### 2.5. Nanoparticle Tracking Analysis (NTA)

Direct visualization, sizing and counting of suspended magnetosome particles was performed using a NanoSight™ LM10 Nanoparticle Analysis System (Malvern Panalytical Ltd, Malvern, Worcs, UK) mounted onto a conventional microscope (fitted with a ×20 objective), equipped with a Marlin F-033B CCD camera (Allied Vision Technologies GmbH, Stadtroda, Germany) for high speed video capture (operated at 30 frames per second), and dedicated Nanoparticle Tracking Analysis software (NTA v1.5).

Following dilution in 0.2 μm filtered PBS and disaggregating using sterile 1 mL hypodermic syringes fitted with ½ inch 29-gauge needles, magnetosome samples (~500 μL) were injected into the sample inlet of the LM10 unit’s 300 μL flow cell. Scattered light from the suspended magnetosome particles interrupting the 80 μm red (635 nm) laser beam passing through the sample chamber was viewed at 20× magnification, and recorded as an 8-bit video file of the particles undergoing Brownian motion (video example in supplementary information, S1). Camera brightness, gain and shutter settings were adjusted in unison to generate optimally exposed blur-free images of polydisperse magnetosome particles (see example screenshot in supplementary information, S2) and a capture duration of 90 s was employed. Image processing was done in expert mode employing 166 nm per pixel, a detection threshold of 33 (Multi), temperature of 22°C, viscosity of 0.9544 cP, with all other analysis parameters set to recommended default values. The first few frames of recorded video were user-adjusted for optimum interference-free tracking in all subsequent frames.

The path taken by each particle in the sample is followed through the video sequence; the image analysis software determining the mean distance moved by each particle in two dimensions (*x* and *y*) over its ‘visible’ lifespan, automatically discounting short-lived and crossing trajectories, and computing its diffusion coefficient, *D_t_*, from the particle’s mean square displacement in two dimensions, 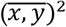. Knowing the sample temperature, *T*, and viscosity, *η*, of the bulk liquid phase the hydrodynamic radius, *r_h_,* is found using Stokes-Einstein equation (Eq. 1) for diffusion in solution, where *K_B_* is Boltzmann’s constant.

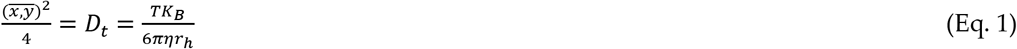

NTA counts particles within a small estimated scattering volume dictated by the optical field of view and depth of the beam and then extrapolates the number seen to an equivalent concentration for particles of a given size and total number of particles, reporting the data as ‘number of particles per mL vs. particle size’ [28, 29]. For deeper analysis of changes in size distribution as functions of ultrasonication time and attack by surfactant the NTA data obtained in this work was minimally is transposed into ‘number’ and ‘number-volume’ based frequency and cumulative undersize distribution plots, from which the corresponding means, medians and modes are found.

### 2.6. Transmission Electron Microscopy (TEM)

Portions (1 mL) of purified magnetosomes suspended in 0.2 μm filtered PBS (prepared as described in section 2.2) were centrifuged at 16,873 g_av_ for 3 min in the FA-45-18-11 fixed angle (45°) rotor of an Eppendorf model 5418 centrifuge (Eppendorf AG, Hamburg, Germany). The pellets were resuspended in 1 mL of 2.5% (v/v) glutaraldehyde in 0.1 M potassium phosphate buffer, pH 7.2, and mixed at 4 rpm for 1 h at room temperature on a TAAB R052 rotator (TAAB Laboratories Equipment Ltd, Aldermaston, Berks, UK). The glutaraldehyde-fixed magnetosome samples were exhaustively dehydrated by a series of washing steps of increasing ethanol concentration (from 50–100% v/v). Sedimented magnetosomes from the last dehydration step were embedded in resin by infiltration of the pellet with a solution containing 50% (v/v) Mollenhauer [39] resin in propylene oxide (Agar Scientific, Stansted, Essex, UK) on a TAAB rotator operated at 4 rpm for 12 h in a fume cupboard, followed by curing in undiluted Mollenhauer resin at 60 °C for another 48 h. Thin sections (120 nm) were cut from the resin block using diamond knives on a Reichert-Jung UltraCut Ultramicrotome (Leica Microsystems GmbH, Wetzlar, Germany). The cut sections were imaged using a JEOL 1200EX transmission electron microscope (JEOL, Tokyo, Japan) operated at 80 keV, in the transmission mode, with the beam current at 60 μA. TEM images were analysed by eye and length of magnetosome chains manually annotated. A minimum of 210 magnetosome crystal units or 62 chains of magnetosomes (whichever was greater) were counted per sample.

## 3. Results and discussion

### 3.1. NTA can be used to quantify magnetosome concentration

The concentration and size distribution of magnetosomes extracted and purified from fed batch grown *M. gryphiswaldense* MSR-1 cells [13] was analysed by NTA. To determine the range of magnetosome concentrations the NTA instrument can be used to accurately measure, dilutions (2, 4, 8, 16, 32 or 64 –fold) of a suspension of ‘batch 1’ magnetosomes (70 mg iron L^−1^) were analysed (Figure 1A). The concentration of the undiluted magnetosome suspension was determined by NTA to be 8.47 × 10^8^ particles mL^−1^. Comparison of the measured magnetosome concentration (Figure 1A, *y*-axis) with the theoretical concentration calculated from the dilution factor (*x*-axis) reveals a linear correlation over the analysed range of 1.3 × 10^7^ and 8.5 × 10^8^ particles per millilitre, in line with previous recommendations for other types of particles [28, 29].

**Figure 1.**
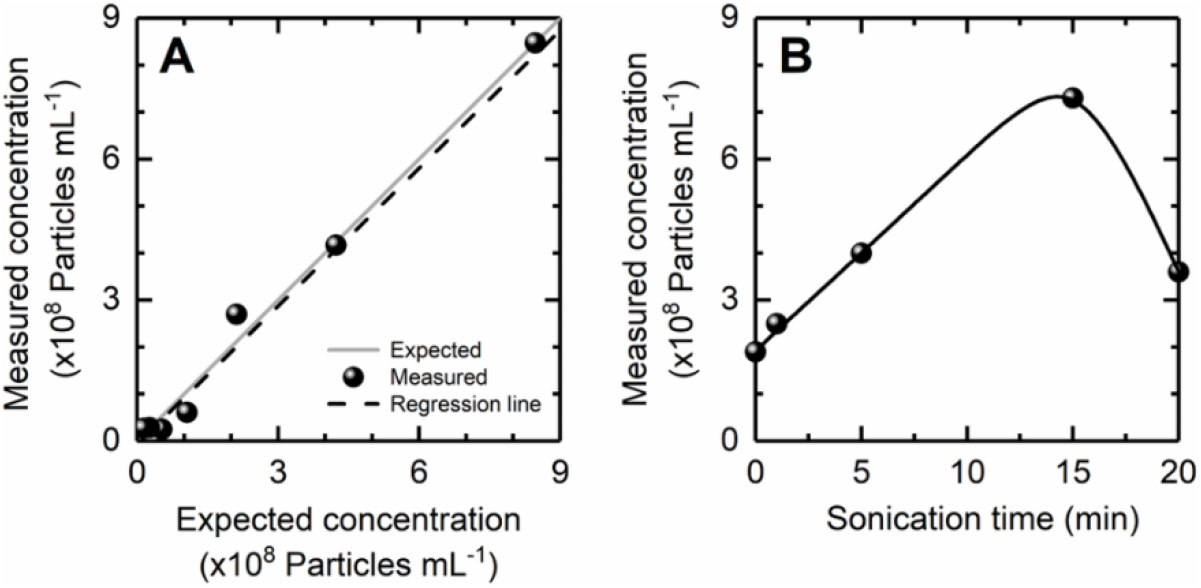
**(A)** Measured vs Expected plot of magnetosome particle concentrations obtained by serial dilution of ‘batch 1’ magnetosomes (70 mg iron L^−1^). A linear regression line (*y* = 1.0346*x* - 0.05602) was fitted to the 2 – 64 diluted sample data points with an R^2^ value of 0.9546. **(B)** Effect of ultrasonication time on magnetosome particle concentration of 4-fold diluted ‘batch 1’ magnetosomes (17.5 mg iron L^−1^).

### 3.2. Effect of ultrasonication time on magnetosome particle concentration and size distribution measured by NTA

Having established the working range of particle concentrations for NTA of magnetosomes, NTA was subsequently employed to study changes in particle distribution and numbers following intentional disruption by liquid shear (ultrasonication) and chemical means (see section 3.3). Ultrasonication has previously been shown to break magnetosome chains [41, 42]; the extent of breakage determined by the operating power and length of exposure. Figure 1B illustrates the effect of ultrasonication time on magnetosome particle concentration. Particle concentration was observe to increase steadily with ultrasonication time, rising from ~2 × 10^8^ particles mL^−1^ initially to a maximum of >7.3 × 10^8^ particles mL^−1^ after 15 min of exposure, before falling steeply to ~3.6 × 10^8^ particle mL^−1^ at the 20 min stage. To understand the reasons for the observed changes in absolute particle numbers we examined the corresponding particle size distributions (Figure 2). In *M. gryphiswaldense* magnetosomes are attached to a protein filament, MamK [40]. Depending on growth conditions each individual *M. gryphiswaldense* magnetosome is 20 – 60 nm in diameter [3 – 6] in chains with average lengths of 14 – 35 magnetosomes [4].

**Figure 2.**
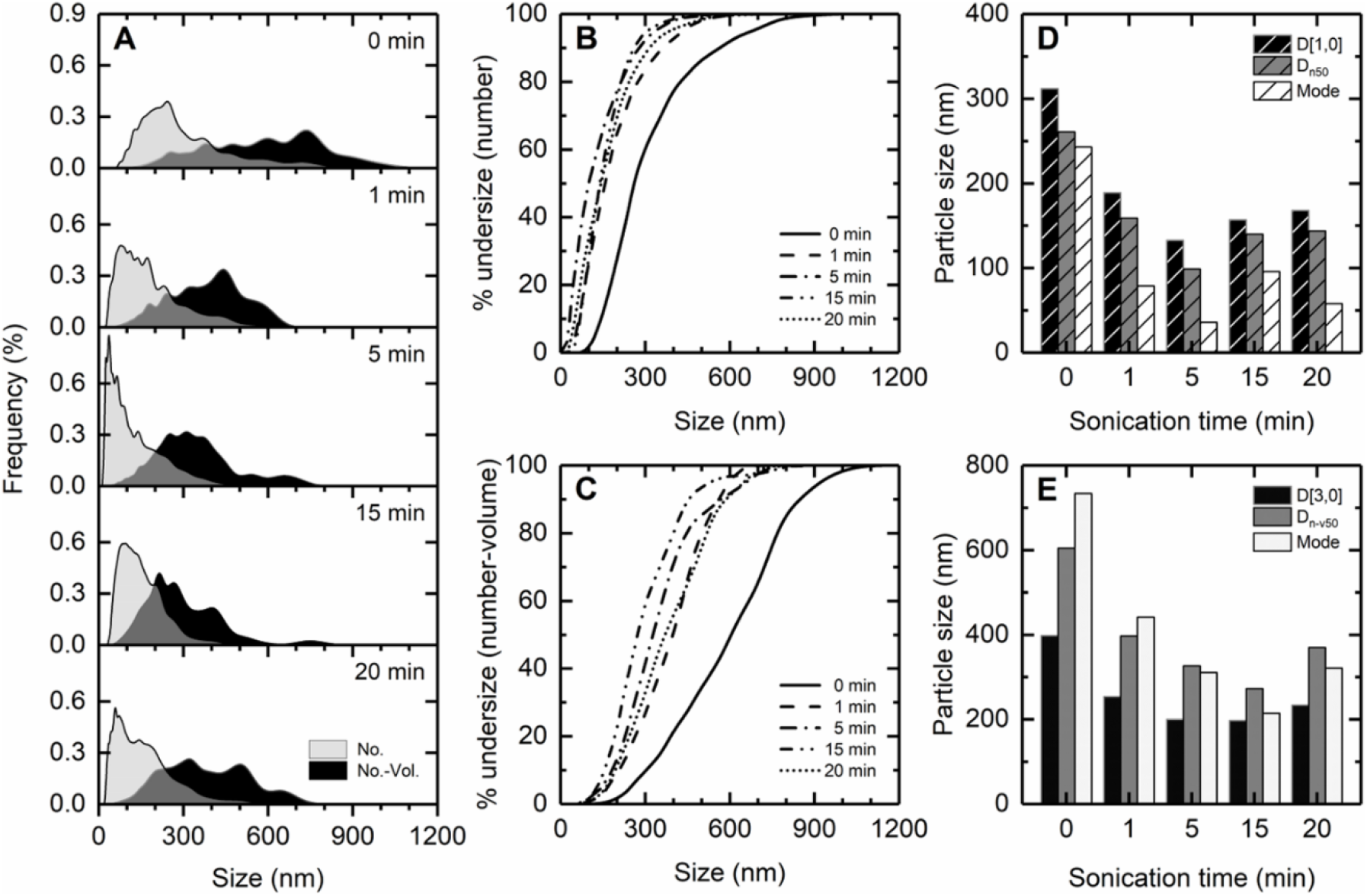
NTA of 4-fold diluted ‘batch 1’ magnetosome suspensions (17.5 mg iron L^−1^) sonicated for different times: **(A)** Number and number-volume frequency distribution plots; **(B)** Number-based cumulative undersize distribution plots; **(C)** D[1,0] mean, D_n50_ and mode values vs. ultrasonication time; **(D)** Number-volume based cumulative undersize distributions plots; **(E)** D[3,0] mean, D_n-v50_ and mode values. The particle concentrations at each time point are given in Figure 1B.

The untreated magnetosome suspension is a polydisperse particle size distribution (Figures 2A – C) characterised by: (i) considerable width (on number and number-volume bases respectively 80% of the magnetosome population, span 143–564 nm and 300–849 nm); (ii) numerous particles of up to 1.1 μm in size; (iii) a number-weighted mean, D[1,0], of 312 nm, median, D_n50,_ of 261 nm, and mode of 243 nm (Figure 2D); and importantly (iv) essentially no particles (<0.002%) in the TEM size range of single isolated magnetosomes, i.e. 40 ± 20 nm (Figures 2A – C). This size distribution strongly infers that the majority of magnetosomes are associated with the MamK fibre, forming chains of various lengths.

Ultrasonication for up to 5 min skewed the distribution profile heavily to smaller sizes (Figure 2A). Clear indicators of increased chain degradation over this time include: (i) increasing numbers of 20 – 60 nm particles appeared in the distribution profiles (accounting for 8.1% of the population after 1 min, rising to 28.2% after 5 min; Figure 2B), giving a number-weighted mode at the 5 min stage of 36 nm, corresponding to individual magnetosomes; and (ii) parallel reductions in D[1,0], D[3,0], D_n50_ and D_n-v50_ values. However, for samples subjected to extended periods of much longer periods of ultrasonication (15 and 20 min) the frequency (Figure 2A) and cumulative distributions (Figures 2B & 2C) profiles shifted back to larger sizes. Very few small particles were observed and all characteristic numbers of the distributions increased *cf.* the 5 min values (Figures 2D & 2E). Taken collectively, the observed changes in overall particle numbers (Figure 1B) and size distribution dynamics (Figure 2) likely reflect two superimposed mechanisms, i.e. fast chain breakage and degradation overlaid by slower aggregation of fragmented magnetosome chains and individual magnetosomes. Agglomeration of magnetosomes is known to be as an adverse consequence of ultrasonication (likely triggered by damage to the magnetosome membrane generating uncoated or partially coated magnetosomes acutely prone to self-aggregation), and the extent to which it occurs increases over time [41, 42]. Comparison of the number-volume frequency distributions between 600 and 800 nm at 1 and 5 min (Figure 2A) reveals agglomeration is already evident at the 5 min stage, but becomes dominant at later time points; corroborated by the dramatic drop in magnetosome concentration from 7.3 × 10^8^ particles mL^−1^ at 15 min to ~3.6 × 10^8^ particles mL^−1^ after 20 min of ultrasonication (Figure 1B).

### 3.3 Effect of SDS on magnetosome particle concentration and size distribution measured by NTA

In a separate experiment, we studied the effect of adding the anionic surfactant, sodium dodecyl sulphate (SDS), to magnetosome suspensions (Figure 3). SDS is commonly employed in the laboratory to disrupt biological membranes, denature and solubilize proteins through concerted interaction of its negatively charged head group and flexible apolar dodecyl tail [43 – 46].

**Figure 3.**
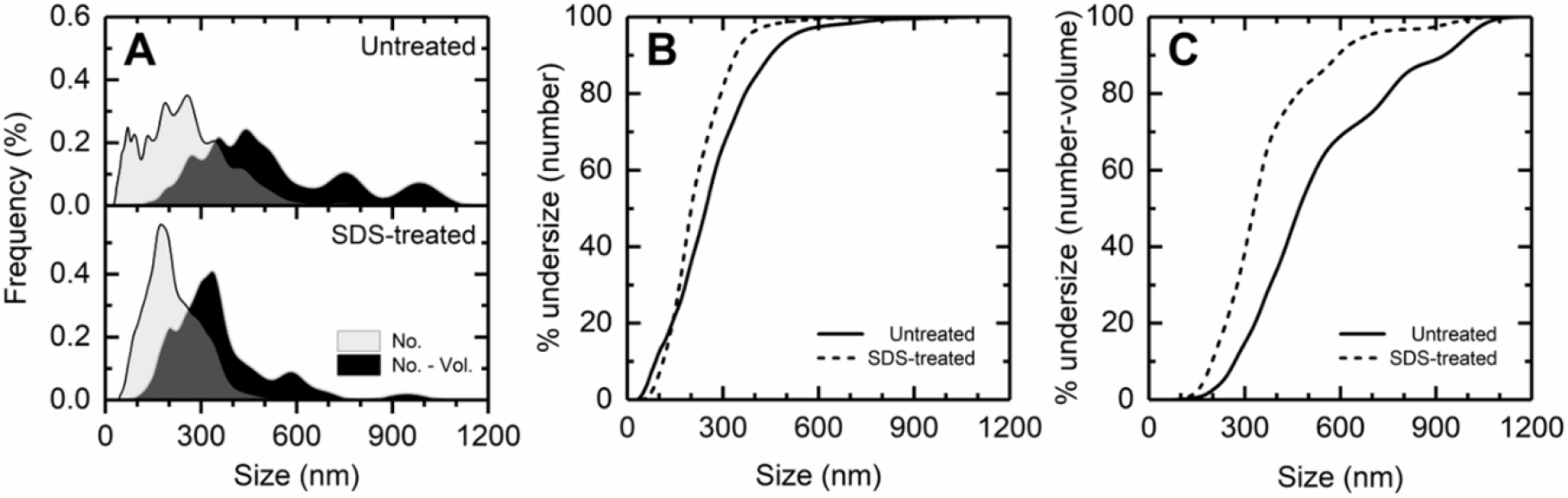
NTA of ‘batch 2’ purified magnetosome (35 mg iron L^−1^) suspensions before and after treatment with 1% (w/v) SDS: (**A)** Number and number-volume frequency distribution plots; **(B)** Number-based cumulative undersize distribution plots; **(C)** Number-volume based cumulative undersize distributions plots. The respective particle concentrations of untreated and SDS-treated ‘batch 2’ magnetosomes were 1.99 × 10^8^ and 5.78 × 10^8^ particles mL^−1^.

Marked changes in both particle number (and distribution profiles were observed following exposure to SDS for 1 h. The particle concentration increased roughly 3-fold from 1.99 × 10^8^ to 5.78 × 10^8^ particles mL^−1^, and all size distributions shifted left to smaller sizes (although the small number of 20–60 nm sized particles in the untreated sample, 3.5%, dropped to 0.4% following SDS treatment) and narrowed considerably. For example, in the case of number based sizing metrics: (i) the arithmetic mean D[1,0] fell from 269 to 219 nm and the standard deviation narrowed from 163 to 102 nm; (ii) D_n50_ decreased from 247 to 201 nm; (iii) mode dropped from 257 to 175 nm; and (iv) 80% of the magnetosome population (i.e. between 10 and 90% cumulative undersize; Figure 3B) was contained within 112 – 339 nm *cf.* 89 – 449 nm for untreated magnetosomes. While inspection of the number-volume frequency (Figure 3A) and cumulative undersize (Figure 3C) plots further emphasise narrowing of the size distributions in the presence *cf.* absence of SDS, with shrinking values for D_n-v10_ (197 *cf.* 269 nm), D_n-v50_ (329 *cf.* 471 nm) and D_n-v90_ (591 *cf.* 926 nm). Note, analogous to D_n-v50_ (the median of the number-volume distribution), D_n-v10_ indicates that 10% has a smaller particle size, and 90% a larger particle size, whereas D_n-v90_ specifies 90% has a smaller particle size, and 10% a larger particle size. The NTA results suggest that SDS may affect magnetosome chains by two main routes, i.e.: (i) denaturation of and/or conferral of negative charge to the MamK polymer filament [40] dissociating magnetosomes from chains; and (ii) damage to the membranes and transmembrane proteins of individual magnetosomes rendering them prone to self-aggregate [41]. The former mechanism affords an explanation for both the loss of larger particles, and 3-fold increase in particle concentration, while the latter explains the loss of the smallest particles from the distribution. When detached from chains individual magnetosomes are much more likely to aggregate into compact structures *cf.* those organised as chains [41, 47].

It can be noted that the size distributions of the untreated magnetosome suspensions depicted in Figures 2 and 3 are quite different. These samples were prepared using different batches of *M. gryphiswaldense* cells, highlighting natural biological variability and reinforcing the requirement for such characterisation methods.

### 3.4. Effect of sonication time on the size distribution of magnetosome examined by TEM

The same sonicated magnetosome suspensions (section 3.2, Figures 1B & 2A – E) were examined by TEM and the resulting micrographs (see Figure 4A for examples) were analysed by eye, recording both the numbers of chains and number of magnetosome units in every chain. For each sonication time point the data is plotted as frequency (number of chains) *vs* number of units per chain (Figure 4B) and % cumulative mass undersize *vs* number of units per chain (Figure 4C). Figure 4D shows the effect of sonication time on mean [D1,0] and median (L_50_) magnetosome chains lengths expressed in magnetosomes per chain.

**Figure 4.**
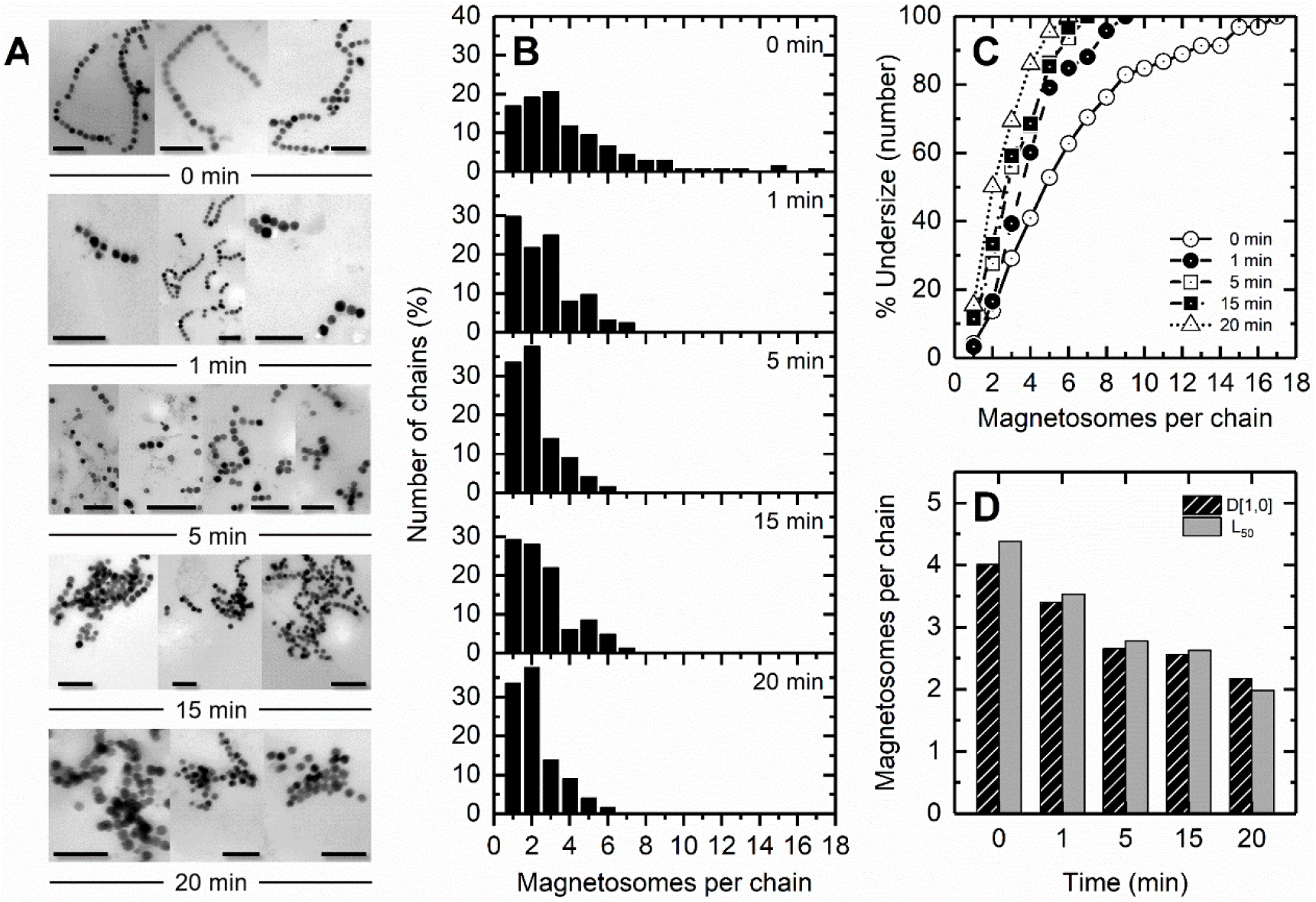
TEM analysis of 4-fold diluted ‘batch 1’magnetosome suspensions (17.5 mg iron L^−1^) sonicated for different times: **(A**) TEM micrographs (Scale bars corresponds to 200 nm); (**B**) Frequency distributions plots; **(C)** mass basis cumulative undersize distribution plots; and **(D)** Dependence of mean [D1,0] and median (L_50_) magnetosome chain length on variation in sonication time. The number of chains (C) and magnetosomes (M) counted at each time point are given in parentheses behind each time point: 0 min (C = 136, M = 545); 1 min (C = 62 C, M = 211); 5 min C = 124, M = 328); 15 min (C = 82, M = 210); 20 min (C = 122, M = 265).

Chains of up to 17 magnetosomes length were observed for the untreated (0 min) sample (Figure 4A & B), but the majority (82.9%) were less than 9 magnetosomes long; the mode, D[1,0] and L_50_ values for the population were respectively 3, 4 and 4.4 magnetosomes per chain. After sonicating for only 1 min chain length was significantly reduced. The small population (17.1%) of longer chains (8 – 17 magnetosomes), being most susceptible to diminution, vanished, and both D[1,0] and L_50_ fell below 3.6 magnetosomes length. With further sonication the distribution profiles gradually shifted to smaller size (Figures 4B and 4C), underlined by parallel incremental declines in D[1,0] and L_50_ at every time point to final estimated lengths of ~2.2 and <2.0 magnetosomes (Figure 4D), although chains of up to 6 magnetosomes still persisted (Figure 4B) albeit in low number (<5% of the population; Figure 4C). For samples exposed to extended sonication, i.e. 15 and 20 min, but not the earlier time points, large dense aggregates of magnetosomes were frequently observed (Figure 4A). As the chain length and number of magnetosomes in such aggregates could not be counted with any degree of certainty our values for D[1,0] and L_50_ values after 15 and especially 20 min of sonication are unlikely to reflect the true distribution of magnetosome sizes in these samples. This said, the propensity for magnetosomes preparations to agglomerate in such a fashion is linked to an increased presence of individual uncoated magnetosome [41, 47]; it follows that the number of single magnetosomes is underestimated and the true mean and median sizes after 20 min of sonication are <2 magnetosomes.

### 3.5. Comparison of NTA and TEM analysis of magnetosome size distribution

Side-by-side comparisons of the frequency distribution profiles measured by NTA (Figure 2A) with those of chain length determined by TEM (Figure 4B) reveal striking similarity during the early time points of ultrasonication (0, 1 and 5 min), with both indicating ~30% of the population (i.e. 29.8% by TEM *cf.* 28.2% by NTA) as single isolated magnetosomes. Comparisons beyond 5 min of ultrasonication are clouded by the TEM analysis, performed here, failing to account for all magnetosome particles in the preparations (see section 3.4). Automated image analysis combined with sophisticated stochastic modelling [26, 48], would ensure greater accuracy in distinguishing and counting magnetosomes in aggregates, but will not resolve all analytical concerns [21]. The complicated sample preparation, potential artefacts introduced and TEM analysis issues encountered here highlight NTA’s utility as complementary low-cost rapid measurement system of particle number and size distribution of magnetosome preparations faithfully reflecting their true state in solution. Several studies, including this one, have shown NTA to be especially useful in detecting and studying aggregation of nanoparticle systems [28, 32 – 36].

## 4. Conclusions

The unique properties of magnetosomes confer key advantages over chemically synthesized magnetic nanoparticles, making them superior prospects for many different applications, especially within the biotech and healthcare sectors [8, 17, 27, 41, 47]. Widespread development of magnetosomes as diagnostic or therapeutic entities will likely hang on parallel advances in enabling technology for future manufacture, formulation and quality control of magnetosome based products [4, 12 – 14]. Key among these will be modern, robust analytical methods for rapid determination of magnetosome concentration, size distribution and aggregation state in solution. Our results demonstrate the feasibility of using NTA for this purpose. However, in its current guise NTA’s ability to resolve magnetosomes of different size and chain length within polydisperse samples is limited at present. Plainly steps to enhance the resolving power of the technique are required. These may include: the use of higher sensitivity cameras and lower wavelength lasers (405, 488 and 532 nm) for more accurate sizing detection of smaller and mid-sized particles in given samples [28, 29, 49]; employing chemically neutral viscosity enhancers to retard particle movement leading to enhanced tracking accuracy [49]; and using improved analytic algorithms that compensate for randomness in Brownian motion to dramatically improve peak isolation and sizing of particles within heterogeneous samples [49]. Work of this nature is currently being done in our laboratories.

## Supplementary Materials

The following are available online

Video S1: Exemplar NTA video of purified magnetosomes.

Figure S1: Exemplar NTA screenshot showing magnetosome chains and individual magnetosomes.

## Acknowledgements

This work was supported by: the ERA-IB grant EIB.13.016 ProSeCa, funded by the UK Biotechnology & Biological Sciences Research Council (BBSRC); an Aston Institute of Materials Research (AIMR) Seed-corn grant; and the EPSRC Centre for Innovative Manufacturing in Emergent Macromolecular Therapies. The authors acknowledge the expert technical assistance of Theresa Morris and Paul Stanley of the University of Birmingham’s Centre for Electron Microscopy.

## Contributions

A.F.C., T.W.O. and O.R.T.T. conceptualized the study; A.F.C., H.L., M.E. and S.J. performed the experiments; M.F., D.G.B., T.W.O. and O.R.T.T. supervised the project; A.F.C., T.W.O. and O.R.T.T. wrote the manuscript; All authors proofread and approved the manuscript.

