## Supplementary figures and images for "Nanoparticle Tracking Analysis: A powerful tool for characterizing magnetosome preparations"

### Figure S1

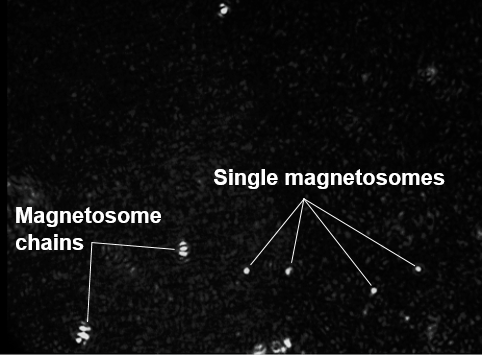
